# Implantation of engineered adipocytes that outcompete tumors for resources suppresses cancer progression

**DOI:** 10.1101/2023.03.28.534564

**Authors:** Hai P. Nguyen, Rory Sheng, Elizabeth Murray, Yusuke Ito, Michael Bruck, Cassidy Biellak, Kelly An, Filipa Lynce, Deborah A. Dillon, Mark Jesus M. Magbanua, Laura A. Huppert, Heinz Hammerlindl, Laura Esserman, Jennifer M. Rosenbluth, Nadav Ahituv

**Affiliations:** Department of Bioengineering and Therapeutic Sciences, University of California San Francisco, San Francisco, CA, USA; Institute for Human Genetics, University of California San Francisco, San Francisco, CA, USA; Division of Hematology/Oncology, Department of Medicine, University of California San Francisco, San Francisco, CA, USA; Dana-Farber Cancer Institute, Harvard University, Boston, MA 02215, USA; Department of Pathology, Brigham and Women’s Hospital, Boston, MA 02115, USA; Department of Laboratory Medicine, University of California, San Francisco, San Francisco, CA 04158, USA; Department of Pharmaceutical Chemistry, University of California, San Francisco, San Francisco, CA 94158, USA; Department of Surgery, University of California, San Francisco, San Francisco, CA 94158, USA; Chan Zuckerberg Biohub – San Francisco, San Francisco, CA 94158, USA

## Abstract

Tumors acquire an increased ability to obtain and metabolize nutrients. Here, we engineered and implanted adipocytes to outcompete tumors for nutrients and show that they can substantially reduce cancer progression. Growing cells or xenografts from several cancers (breast, colon, pancreas, prostate) alongside engineered human adipocytes or adipose organoids significantly suppresses cancer progression and reduces hypoxia and angiogenesis. Transplanting modulated adipocyte organoids in pancreatic or breast cancer mouse models nearby or distal from the tumor significantly suppresses its growth. To further showcase therapeutic potential, we demonstrate that co-culturing tumor organoids derived from human breast cancers with engineered patient-derived adipocytes significantly reduces cancer growth. Combined, our results introduce a novel cancer therapeutic approach, termed adipose modulation transplantation (AMT), that can be utilized for a broad range of cancers.

## Introduction

Tumors are complex tissues composed of cancerous and non-cancerous cells in a hypoxic and nutrient-deprived microenvironment. The tumor microenvironment (TME) contains heterogeneous cell populations, including immune cells, mesenchymal support cells, and matrix components contributing to tumor growth and progression (*1*). In order to survive this environment, tumors are capable of reprogramming metabolic pathways to better utilize available substrates in the surrounding TME, ultimately becoming dependent on these pathways for continued growth and survival (*2*). In contrast to normal cells, the main pathway of glucose metabolism in cancer cells is aerobic glycolysis, termed the Warburg effect (*3*). Glucose uptake and production of lactate is increased in these cells, even in the presence of oxygen and functional mitochondria (*3*). The increase in glycolytic flux allows glycolytic intermediates to supply subsidiary pathways to fulfill the metabolic demands of proliferating cells. During hypoxia, cancer cells also undergo metabolic reprogramming to increase lipid utilization, as fatty acids produce twice the energy of glucose (*4, 5*).

There are many efforts to target cancer glucose and fatty acid metabolism for therapeutic purposes. For glycolysis, these include drugs that target hexokinase 2 (HK2), which is involved in the initial steps in glycolysis, using ATP from the mitochondria to phosphorylate glucose, or drugs that target glucose transporters (GLUT1 and GLUT4)(*2, 6-8*). Several drugs are also used to target lipid metabolism in cancer(*9-12*). These include drugs that target lipid uptake (targeting proteins such as LXR, CD36 and FABP4/5), lipogenic enzymes (such as *ACC, ACLY, FASN*, and *SCD1*) and proteins involved in intracellular lipid homeostasis (such as *CPT1A, PPARG* and others)(*13, 14*). In addition, recent work has shown that cold activation of brown adipose tissue (BAT), which dissipates energy by non-shivering thermogenesis, increases adipocyte glucose uptake, lipid metabolism and significantly inhibits tumor growth (*15*). However, situating cancer patients in cold conditions for extended periods is challenging.

Here, we set out to develop a novel cancer therapeutic approach, termed adipose manipulation transplantation (AMT) that utilizes two unique abilities of white adipose tissue (WAT): 1) They can be readily extracted via liposuction and implanted via reconstructive surgery; 2) they can change into a BAT-like tissue, called browning or beiging (*16, 17*), by upregulating essential transcriptional regulators or enzymes, such as the uncoupling protein 1 (*UCP1*), PPARG coactivator 1 alpha (*PPARGC1A*) or PR/SET domain 16 (*PRDM16*), involved in BAT development and function (*18-33*). Similar to BAT, beige adipocytes have the capacity to convert energy to heat and contribute to whole-body energy expenditure (*34*). We show that CRISPR activation (CRISPRa) of either *UCP1, PRDM16* or *PPARGC1A*, induces browning which subsequently increases glucose and fat metabolism in human white adipocytes and adipocyte organoids. Co-culturing of these CRISPRa-modulated adipocytes with various cancer cell lines (breast, colon, pancreatic, or prostate cancer) leads to significantly suppressed cancer cell proliferation as well as decreased glucose uptake, glycolysis and fatty acid oxidation capacity in the cancer cells. Subcutaneously co-transplanting CRISPRa-modulated human adipose organoids and cancer cells (breast, colon, pancreatic, or prostate) in immune-compromised mice leads to significantly reduced tumor size with decreased hypoxia and angiogenesis. Implantation of engineered adipose organoids in pancreatic or breast cancer genetic mouse models significantly reduces cancer progression. Furthermore, in the breast cancer model we show that their implantation both near and distal to the tumor leads to similar results. To further demonstrate the therapeutic potential of this approach, we show that adipocytes isolated from human dissected breast tissues can be similarly manipulated via CRISPRa and inhibit growth of patient-derived breast cancer organoids. Combined, our results introduce a novel cancer therapeutic approach that has the potential to treat numerous cancer types.

## Results

### CRISPRa browning of human white adipocytes

To induce browning in human adipocytes we utilized CRISPRa to upregulate *UCP1, PPARGC1A* or *PRDM16*, all known genes involved in BAT development and function. Using CRISPick (*35*), we designed five gRNAs targeting each gene’s promoter and cloned them into an adeno associated virus (AAV) based expression vector.

Differentiated adipocytes derived from human white preadipocytes were co-transfected with the gRNAs along with a *Staphylococcus aureus* (sa) endonuclease deficient Cas9 (dCas9) fused to the VP64 transcriptional activator. We used an sa dCas9 due to its smaller size and the VP64 transcriptional activator, which carries four copies of VP16, a herpes simplex virus type 1 transcriptional activator (*36*), as it provides moderate gene upregulation and is small enough to fit into AAV, which has a 4.7 kilobase optimal packaging capacity (*37*). To generate mature human adipocytes, preadipocytes were subjected to adipocyte differentiation using a cocktail of 3-isobutyl-1-methylxanthine (IBMX), dexamethasone, and insulin before being subjected to CRISPRa. Following four days, we measured gene expression via qRT-PCR, finding several gRNAs to significantly increase the expression levels of *UCP1, PPARGC1A, PRDM16* in the transfected cells compared to the control cells (treated with dCas9-VP64 only) (**fig.S1A**). We selected the top two gRNA for each gene for AAV serotype 9 (AAV9) packaging. We used AAV9, as it was shown to effectively infect various adipose depots (*38*). We infected human differentiated adipocytes with these viruses along with AAV9 dCas9-VP64, finding for each gene at least one gRNA that significantly increased expression levels compared to dCas9-VP64 only control **(fig.S1B**). We used the top activating gRNA AAV for all subsequent experiments.

We next examined whether our CRISPRa treatment increases browning in these human white adipocytes. Human adipocytes infected with the top gRNA for *UCP1, PRDM16* or *PPARGC1A* showed significant upregulation of their target genes compared to dCas9-VP64 only infected cells (**Fig.1A**). In addition, we also observed increased mRNA levels for brown fat marker genes, including *TFAM, DIO2, CTP1b*, and *NRF1* upon upregulation of either of the three genes (**Fig.1B**; *PRDM16* CRISPRa did not show upregulation of *CTP1b*, and *NRF1*). We examined oxygen consumption rate (OCR) in these cells using Seahorse (see Methods), by initially blocking oxygen consumption and then adding 1 μM oligomycin, followed by the introduction of 1 μM FCCP to measure the maximal respiratory capacity. We found that CRISPRa-AAV treatment targeting either gene increased overall OCR levels in human white adipocytes with *UCP1* gRNA AAV treated cells having the largest increase (**Fig.1C**). In addition, these cells showed increased uncoupled respiration (under oligomycin, ATPase inhibitor) indicating a brown fat-like phenotype (**Fig.1C**). CRISPRa treated cells also had elevated maximal respiration following carbonyl cyanide-p-trifluoromethoxy-phenylhydrazone (FCCP) treatment (**Fig.1C**). Furthermore, these AAV-CRISPRa engineering adipocytes showed increased glucose uptake in both basal and insulin stimulated conditions (**Fig.1D**). We also tested the fatty acid oxidation (FAO) capacity of these cells by performing a similar Seahorse OCR assay. In both BSA- and BSA conjugated-palmitate (saturated fatty acid complex) media, we found that our CRISPRa-treated adipocytes had an overall OCR increase in BSA conjugated-palmitate (**Fig.1E**). Under FCCP treatment, in which the increase of OCR in palmitate-containing media compared to BSA-containing media is thought to be due to exogenous fatty acid oxidation, we found that upregulation of either gene increased exogenous FAO capacity in human adipocytes (**Fig.1F**). Taken together, we show that AAV-based CRISPRa upregulation of *UCP1, PPARGC1A* or *PRDM16* induces browning in human adipocytes, leading them to have increased glucose uptake and fatty acid oxidation.

**Fig. 1.**
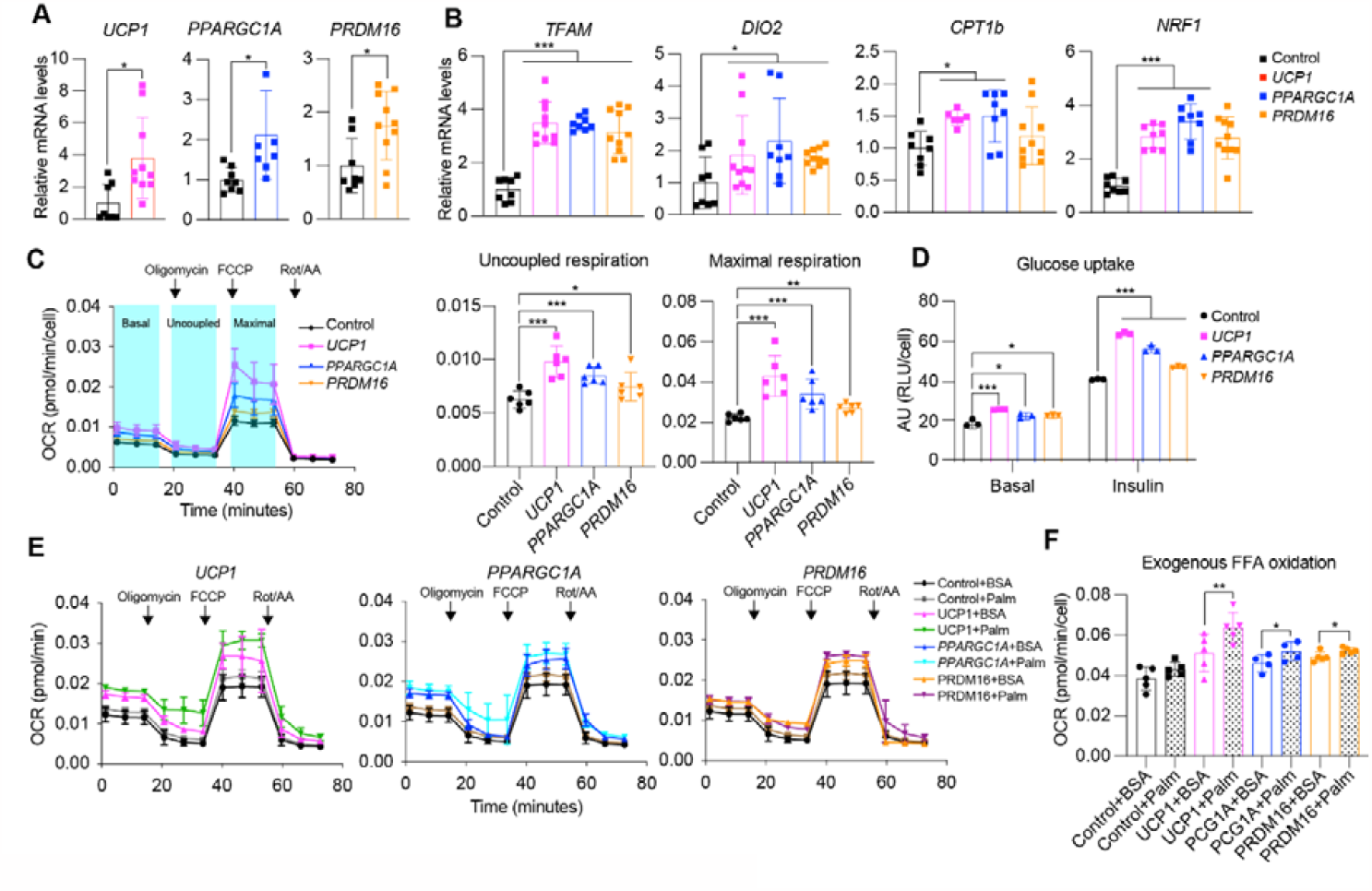
CRISPRa browning activation in human white adipocytes. (**A**) qRT-PCR of *UCP1, PPARGC1A*, and *PRDM16* in human white adipocytes transduced with CRISPRa targeting *UCP1, PPARGC1A*, and *PRDM16*. Data are represented as mean ± S.D *≤0.05. (**B**) qRT-PCR of *TFAM, DIO2, CPT1b*, and *NRF1* in CRISPRa-modulated adipocytes. Data are represented as mean ± S.D *≤0.05, ***≤0.001. (**C**) Oxygen consumption rate (OCR) of CRISPRa-modulated adipocytes measured by the seahorse assay. Uncoupled OCR was measured under oligomycin treatment, while maximal OCR was measured under FCCP. (**D**) Glucose uptake of CRISPRa-modulated adipocytes with or without insulin. Data are represented as mean ± S.D *≤0.05, **≤0.01, ***≤0.001. (**E**) OCR of CRISPRa-modulated cells was measured by the seahorse assay in BSA- or BSA-Palmitate-medium. Data are represented as mean ± S.D *≤0.05, ***≤0.001. (**F**) Exogenous fatty acid oxidation of CRISPRa-modulated adipocytes was calculated by the difference of area under the curve of OCR between BSA- and BSA-Palmitate-media upon FCCP treatment. Data are represented as mean ± S.D *≤0.05, **≤0.01.

### CRISPRa-modulated adipocytes suppress cancer cell growth *in vitro*

We next evaluated whether our CRISPRa ‘browned’ adipocytes could inhibit cancer growth *in vitro* using a co-culturing system (**Fig.2A**). We initially treated differentiated adipocytes with CRISPRa-AAV for either of the three genes (*UCP1, PPARGC1A* or *PRDM16*) and replated these adipocytes on the top chamber of a 12- or 24-well transwell plate, which has inserts of 0.4um membrane, so that the adipocytes don’t contact the cells on the lower chamber (**Fig.2A**). In the lower chamber, we grew five different cancer cell lines: breast cancer cells MCF7 (ER+,PR+,GR+) and MDA-MB-436 (triple negative), colon cancer (SW-1417), pancreatic cancer (Panc 10.05) and prostate cancer (DU-145). As a negative control, we used adipocytes infected with dCas9-VP64 only. After three days, we observed that all five cancer cell lines that were co-cultured with *UCP1, PPARGC1A* or *PRDM16* CRISPRa-treated human adipocytes showed significantly lower cell numbers than cells co-cultured with dCas9-VP64 treated adipocytes (**Fig.2B**). We found that the number of cancer cells co-cultured with CRISPRa-treated adipocytes were 3- to 5-fold lower than cells cocultured with the control adipocytes (**Fig.2C**). By qRT-PCR, we observed that all cancer cells co-cultured with CRISPRa-treated adipocytes had significantly decreased levels of *MKI67* (other than DU-145 treated with *PRDM16* CRISPRa) a proliferation marker, compared to control cells, with *UCP1* CRISPRa having the greatest effect (**Fig.2D**). We also performed a BrdU incorporation assay, finding that after 24 hours the CRISPRa co-cultured cancer cells showed significantly reduced proliferation compared to cells co-cultured with control adipocytes (**fig.S2A**).

**Fig. 2.**
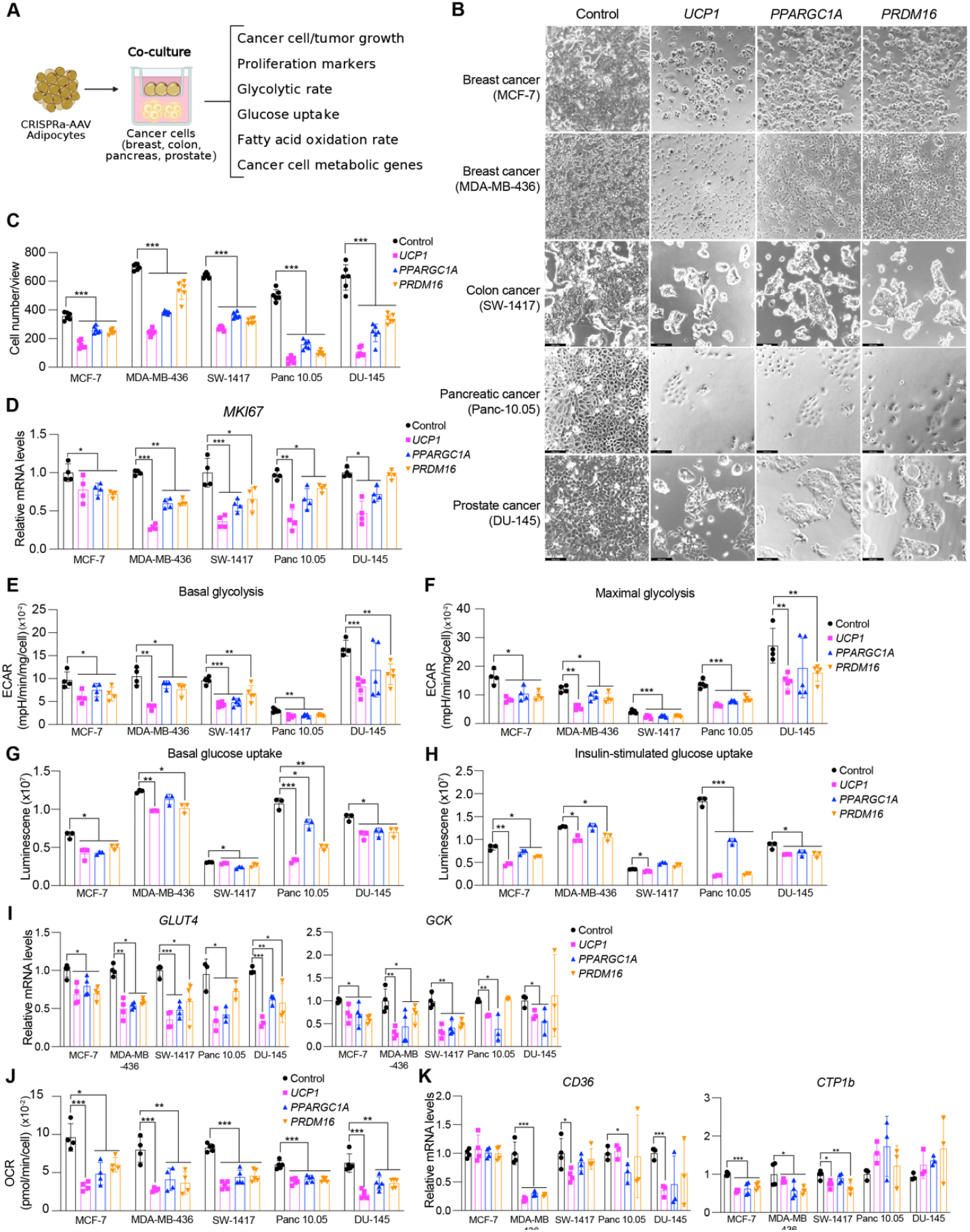
CRISPRa-modulated adipocytes inhibit cancer cell growth *in vitro*. (**A**) Schematic of the co-culturing model of cancer cells and CRISPRa-treated adipocytes using transwell plates and their subsequent phenotyping. (**B**) Representative images of cancer cells, including breast (MCF7, MDA-MD-436), colon (SW-1417), pancreatic (Panc 10.05), and prostate cancer (DU-145) that were co-cultured with CRISPRa upregulating *UCP1, PPARGC1a*, and *PRDM16* or control (dCas9-VP64 only) adipocytes. (**C**) Cancer cell numbers per view of image (4 images/replicates per condition). Data are represented as mean ± S.D ***≤0.001. (**D**) qRT-PCR of the proliferation marker gene, *MKI67*, for cancer cells co-cultured with CRISPRa-modulated adipocytes. Data are represented as mean ± S.D *≤0.05, **≤0.01, ***≤0.001. (**E**) Basal glycolysis measured by calculating area under the curve of extracellular acidification rate upon glucose treatment. Data are represented as mean ± S.D *≤0.05, **≤0.01, ***≤0.001. (**F**) Maximal glycolysis measured by calculating area under the curve of extracellular acidification rate upon oligomycin treatment. Data are represented as mean ± S.D *≤0.05, **≤0.01, ***≤0.001. (**G**-**H**) Glucose uptake of cancer cells co-cultured with CRISPRa-modulated adipocytes without (**G**) or with (**H**) insulin. Data are represented as mean ± S.D *≤0.05, **≤0.01, ***≤0.001. (**I**) qRT-PCR of glucose transporter, *GLUT4*, and glycolytic enzyme, *GCK* in cancer cells. Data are represented as mean ± S.D *≤0.05, **≤0.01, ***≤0.001. (**J**) Exogenous fatty acid oxidation of cancer cells calculated by the difference of area under the curve of OCR between BSA- and BSA-Palmitate-media upon FCCP treatment. Data are represented as mean ± S.D **≤0.01, ***≤0.001. (**K**) qRT-PCR of fatty acid transporter, *CD36*, and fatty acid regulatory transporter, *CPT1b* in cancer cells that were co-cultured with CRISPRa-treated adipocytes. Data are represented as mean ± S.D *≤0.05, ***≤0.001.

We next analyzed the glucose and fatty acid metabolism of the cancer cells. To measure glycolysis, we utilized the extracellular acidification rate (ECAR) assay in which the basal glycolytic rate is measured by adding glucose and the maximal glycolytic rate is measured by oligomycin A addition. We found that all five cancer cell lines co-cultured with CRISPRa-AAV treated adipocytes showed lower overall glycolysis than those co-cultured with dCas9-VP64 only (**fig.S2B**). These cells also showed a significant reduction in both basal and maximal glycolytic rate (**Fig.2E-F**) and lower glucose uptake in both basal and insulin conditions (**Fig.2G-H**), other than SW-1417 treated with either *PPARGC1A* or *PRDM16* CRISPRa. Using qRT-PCR, we also found that the expression of key glycolysis genes, such as *GCK* and *GLUT4*, a major glucose transporter, were significantly lower in the cancer cells co-cultured with CRISPRa-AAV adipocytes (other than *GCK* for *PRDM16* CRISPRa treated Panc 10.05 and DU-145 cells) compared to the negative control (**Fig.2I**). We also examined fatty acid oxidation (FAO) using Seahorse. Similar to glycolysis, we found that all five cancer cell lines that were co-cultured with CRISPRa-AAV adipocytes had reduced FAO compared to the negative control (**Fig.2J, fig.S2C**). Moreover, using qRT-PCR, we found in most CRISPRa conditions that the cancer cells have lower expression of both *CPT1b*, a key regulator of FAO in the mitochondria and *CD36*, a fatty acid transporter on the cell membrane, further confirming decreased FAO (**Fig.2J**). Combined, our data shows that our CRISPRa-modulated adipocytes reduce glycolysis and fatty acid metabolism in five different cancer cell lines and can significantly suppress cancer growth.

### CRISPRa-modulated human adipose organoids suppress xenograft growth

To examine whether CRISPRa-modulated adipocytes could inhibit cancer growth in *in vivo* xenograft models, we co-transplanted four cancer cell lines (MCF7 and MDA-MB-436 (breast), Panc 10.05 (pancreas) and DU-145 (prostate)) with CRISPRa-modulated human adipose organoids. While adipocytes could be used for co-transplantation, adipose organoids offer added advantages, including: 1) providing a 3D-culture that can better recapitulate the heterogeneity of adipose tissue; 2) enhanced response to endogenous stimuli; 3) the ability to form tissue microenvironments that could better integrate with cancer cells following transplantation. We initially established culturing conditions for human adipose organoids. We utilized immortalized human preadipocytes grown using methods that combines features from three different 3D culturing protocols (*39, 40*). In brief, we cultured human preadipocytes in basal DMEM media supplemented with 10% fetal bovine serum in Nunc 96 well plates treated with Nunclon Delta. Organoids formed after 48 hours and were then differentiated to adipose organoids using a differentiation cocktail containing IBMX, dexamethasone, insulin, T3, and rosiglitazone. Adipocytes formed 21-days post differentiation (**fig.S3A**). Using qRT-PCR, we analyzed these organoids for various adipogenic markers, including *FABP4, PLIN1*, and *ADIPOQ*, finding all to be expressed (**fig.S3B**). We further tested our ability to upregulate *UCP1, PPARGC1A*, and *PRDM16* in these organoids using similar methods and AAV gRNAs as in the adipocytes, finding all three to show significant upregulation of the target genes (**fig.S3C**). With the establishment of these adipose organoid culturing and CRISPRa conditions, we next set out to test whether they can suppress xenograft cancer growth.

To generate xenografts, cancer cells were subcutaneously implanted into immuno-compromised SCID mice. Following 6-8 weeks, *UCP1*-CRISPRa treated human adipose organoids were mixed with Matrigel and co-transplanted adjacent to palpable tumors. For all subsequent assays, we only used *UCP1*-CRISPRa, as it showed the most optimal results in our cell culture experiments. Due to some reports suggesting that brown fat could be linked to cancer-associated cachexia (loss of skeletal muscle and fat) (*41*), we measured on a weekly basis the body weight of mice co-transplanted with cancer cells and *UCP1*-modulated adipose organoids, finding no significant differences between CRISPRa treated and control mice (**fig.S4D**). Tumors and human adipose organoids were collected following three weeks (**Fig.3A**). All tumor types co-transplanted with CRISPRa-modulated human adipose organoids were significantly smaller than dCas9-VP64 only transplanted human adipose organoids (**Fig.3B**), having over 50% reduction in volume (**Fig.3C**). Gene analysis showed that tumors co-transplanted with CRISPRa-treated adipose organoids had decreased expression of the proliferation marker gene *MIK67* (**fig.S3E**), as well as reduced marker gene expression for glycolysis (*GLUT4, GCK*) and fatty acid oxidation (*CD36, CPT1B*) (**fig.S3F**). Using immunofluorescence, we examined additional tumor marker genes, finding that all tumors co-transplanted with *UCP1*-CRISPRa modulated human adipose organoids had markedly reduced Ki67+ (**Fig.3D**). In addition, cancer cells showed decreased levels of hypoxia, identified by having a lower carbonic anhydrase (CA9+) area per image view (**Fig.3E**). These tumors also showed decreased levels of CD31+ area per image view, indicating decreased microvessel density and suggesting a corresponding lower metastatic potential (**Fig.3F)**. Since brown fat is known to reduce insulin levels, we examined the plasma insulin in these mice. We found that plasma insulin levels of mice that were co-transplanted with *UCP1*-CRISPRa adipose organoids and cancers were significantly lower than those co-transplanted with the control adipose organoids and were comparable to levels in wild type SCID mice (**fig.S3G**). Taken together, these results show that *UCP1*-CRISPRa modulated human adipose organoids significantly reduce glycolysis and fatty acid oxidation, reduce hypoxia and insulin levels and inhibit tumor growth for four different cancer types *in vivo*.

**Fig. 3.**
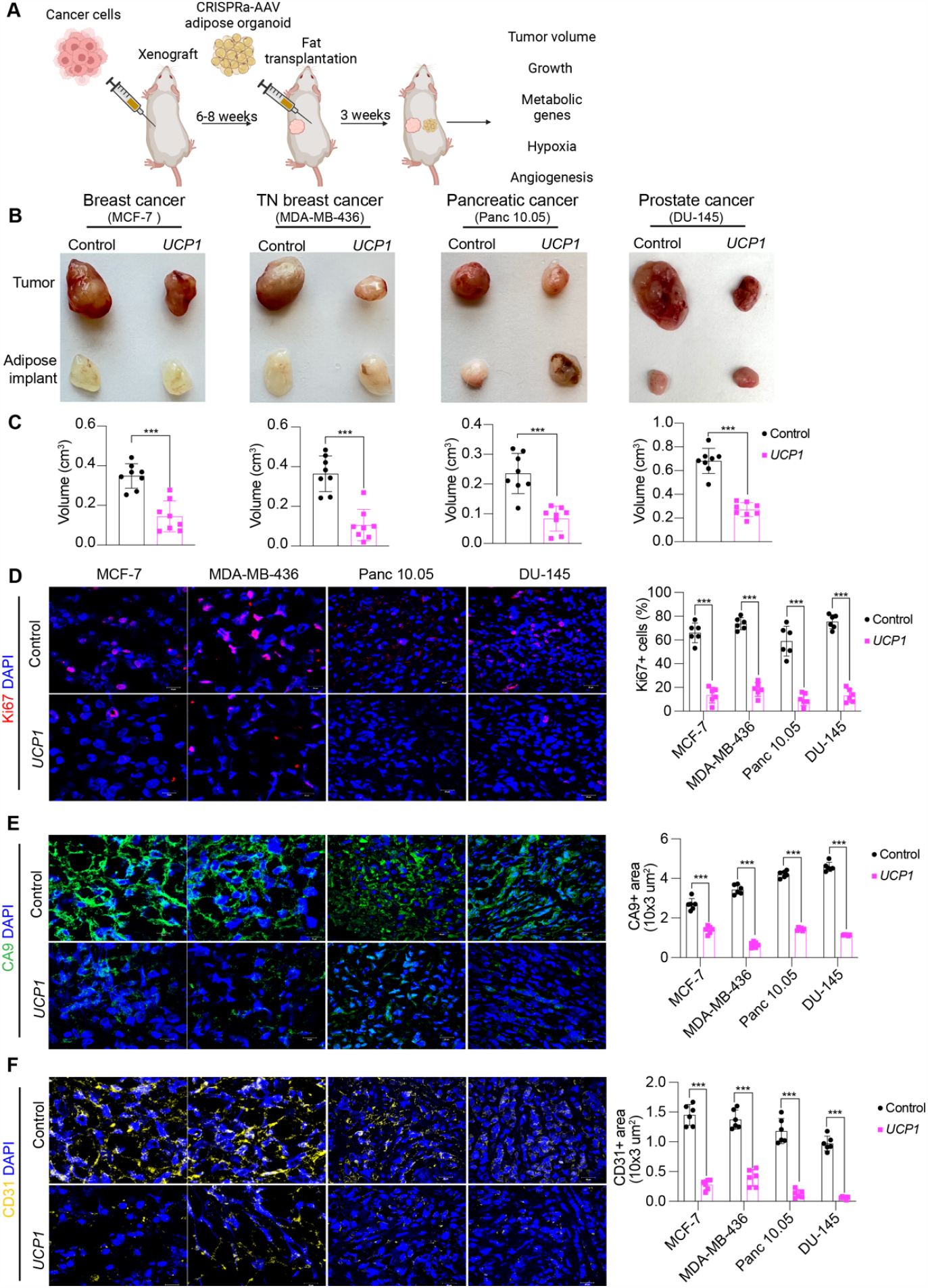
Co-transplantation of xenografts with *UCP1*-CRISPRa modulated human adipose organoids suppresses tumor growth. (**A**) Schematic of the co-transplantation model for xenografts and *UCP1*-CRISPRa treated human adipose organoids in immune-deficient SCID mice and their subsequent phenotyping. (**B**) Representative images of xenograft tumors from various cancer cells lines, including breast (MCF7 and MDA-MD-436), pancreatic (Panc 10.05), and prostate cancer (DU-145) that were co-transplanted with *UCP1*-CRISPRa human adipose organoids or control (dCas9-VP64 only) adipose organoids (n=8 mice per treatment). (**C**) Volume of xenograft tumors that were co-transplanted with *UCP1*-CRISPRa human adipose organoids compared to control (dCas9-VP64 only) (n=6-8 mice). Data are represented as mean ± S.D ***≤0.001. (**D**-**F**) Immunofluorescence staining and quantification of Ki67 (**D**), CA9 (**E**), and CD31 (**F**) in cryosections of xenograft tumors (n= 4 sections per treatment). Data are represented as mean ± S.D ***≤0.001.

### AMT suppresses cancer development in pancreatic and breast cancer genetic mouse models

To examine whether our AMT approach can prevent cancer development, we utilized pancreatic and breast cancer genetic mouse models. For pancreatic cancer, we used the KPC mouse model, that upon tamoxifen treatment develops pancreatic ductal adenocarcinoma due to conditional mutations in *Kras* and *Trp53* (42). For breast cancer, we used *MMTV-PyMT* mice on an FVB background, containing a mouse mammary tumor virus (MMTV) long terminal repeat upstream of the Polyoma Virus middle T antigen (PyVmT), that develop mammary tumors with a mean latency of 53 days (*43*). We first designed five gRNAs targeting mouse *Ucp1* and transfected them into mouse 3T3-L1 adipocytes, finding all five upregulate *Ucp1* (**fig.S4A**). We next selected the top two gRNA for AAV serotype 9 (AAV9) packaging and infected 3T3-L1 differentiated adipocytes with these viruses along with AAV9 dCas9-VP64, finding that all of them significantly increased *Ucp1* gene expression levels compared to dCas9-VP64 only control (**fig.S4B**). We used the top activating gRNA AAV9 for all subsequent experiments. As organoids have several of the aforementioned advantages, we generated adipose organoids using mouse preadipocytes, utilizing similar techniques as described for human adipose organoids (see Methods). We then infected them with AAV9 dCas9-VP64 and *Ucp1*-gRNA and observed both *mCherry* expression from our gRNA virus and significant *Ucp1* upregulation (**fig.S4C**).

We first set out to test whether we can suppress pancreatic cancer development using KPC mice. KPC mice were treated with tamoxifen postnatal day 0-4. At four weeks of age, *Ucp1*-CRISPRa or dCas9-VP64 only (negative control) adipose organoids were orthotopically implanted next to the pancreas (**Fig.4A**). Following six weeks, pancreases were removed and analyzed. Over these six weeks, we observed no significant difference in body weight between mice implanted with *Ucp1*-CRISPRa organoids or control organoids (dCas9-VP64 only) (**fig.S4D**). We found that KPC mice implanted with *Ucp1*-CRISPRa upregulated adipose organoids had significantly smaller tumors compared to control dCas9-VP64 implanted mice (**Fig.4B**) and reduced pancreatic mass (**Fig.4C**). They also showed reduced expression of the proliferation marker, *Mki67*, as well as genes involved fatty acid oxidation (*Cd36, Cpt1b*) (**Fig.4D**). In addition, we found *Ucp1*-CRISPRa mice to have lower expression levels of the pancreas’ main glucose transporter, *Glut2* (**Fig.4D**). *Ucp1*-CRISPRa mice also had a lower number of Ki67+ cells compared to control mice as determined by immunofluorescence (**Fig.4E, S4E**). Similarly, we observed lower CA9+ area per image, and CD31+ area per image view, suggesting reduced hypoxia and angiogenesis (**Fig.4E, S4E**). In addition, we found *Ucp1*-CRISPRa adipose organoids to have significantly reduced insulin levels compared to control mice (**fig.S4F**).

**Fig. 4.**
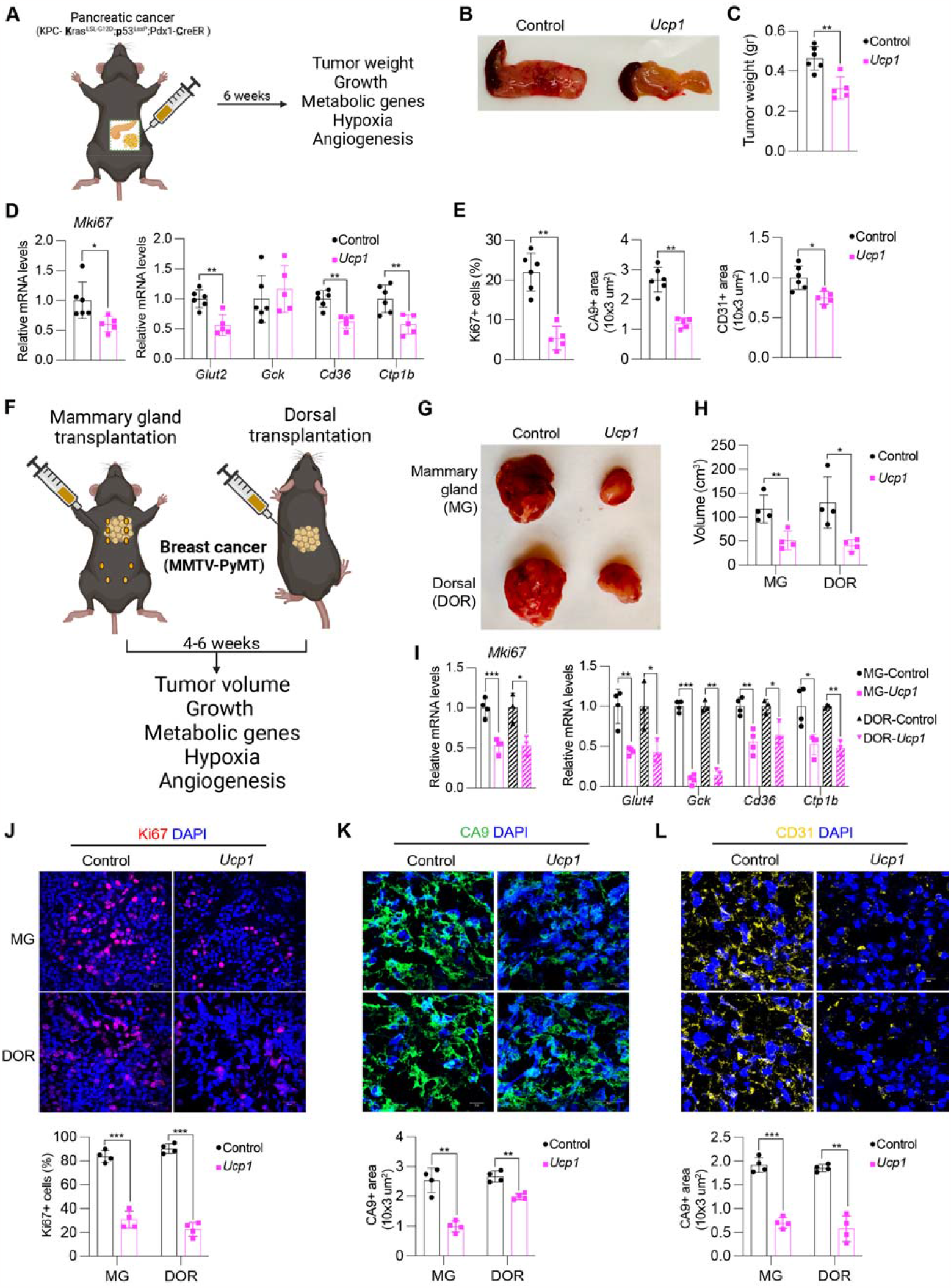
Implantation of *Ucp1*-CRISPRa adipose organoids in pancreatic and breast cancer genetic mouse models suppresses cancer development. (**A**) Schematic of the transplantation model for *Ucp1*-CRISPRa treated mouse adipose organoids in KPC pancreatic cancer mice and their subsequent phenotyping. (**B**) Representative images of the pancreas implanted with *Ucp1*-CRISPRa or control (dCas9-VP64 only) mouse adipose organoids (n=5-6 mice per treatment). (**C**) Weight of the pancreas transplanted with *Ucp1*-CRISPRa modulated mouse adipose organoids compared to control (dCas9-VP64 only) (n=5-6 mice). Data are represented as mean ± S.D **≤0.01. (**D**) qRT-PCR of proliferation marker gene, *MKI67*, metabolic genes, including *Glut2, Gck, Cd36*, and *Ctp1b*, from pancreatic tumors co-transplanted with *Ucp1*-CRISPRa modulated adipocytes. Data are represented as mean ± S.D *≤0.05, **≤0.01. (**E**) Immunofluorescence quantification of Ki67, CA9, and CD31 in cryosections of tumors (n= 4 sections per treatment). Data are represented as mean ± S.D *≤0.05, **≤0.01. (**F**) Schematic of the transplantation model for *Ucp1*-CRISPRa treated mouse adipose organoids in the mammary gland or on the back of *MMTV-PyMT* breast cancer mice and their subsequent phenotyping. (**G**) Representative images of the breast tumor that were implanted with *Ucp1*-CRISPRa or control (dCas9-VP64 only) adipose organoids in the mammary gland or on the back of the mice (Dorsal) (n=4-5 mice per treatment). (**H**) Volume of the tumors transplanted with *Ucp1*-CRISPRa adipose organoids compared to control (dCas9-VP64 only) (n=4-5 mice). Data are represented as mean ± S.D *≤0.05, **≤0.01. (**I**) qRT-PCR of proliferation marker gene, *MKI67*, metabolic genes, including *GLUT4, GCK, CD36*, and *CPT1b*, from breast tumors co-transplanted with *Ucp1*-CRISPRa modulated adipocytes. Data are represented as mean ± S.D *≤0.05, **≤0.01, ***≤0.001. (**J-L**) Immunofluorescence staining and quantification of Ki67 (**J**), CA9 (K), and CD31 (**L**) in tumor cryosections (n= 4 sections per treatment). Data are represented as mean ± S.D **≤0.01, ***≤0.001.

To examine whether our treatment might provide systematic therapeutic effect in suppressing breast cancer growth, we next used *MMTV-PyMT* female mice, implanting *Ucp1*-CRISPRa or dCas9-VP64 adipose organoids near the third nipples (MG) of four-week-old mice (**Fig.4F**). As cold treatment leads to widespread BAT activation causing cancer suppression (*15*), we also examined whether distal implantation of adipose organoids could suppress cancer development by implanting organoids in the back of 4-week-old mice (DOR). Following six weeks post implantation, tumors were collected and analyzed. Similarly, we found no difference in body weight of mice implanted with *Ucp1*-CRIPSRa organoids compared to those implanted with control organoids (**fig.S4G**). Remarkably, we found both strategies of organoid implantation resulted in significantly reduced tumor size (**Fig.4G**) and volume (**Fig.4H**) regardless of site of implantation. Tumors also had decreased expression of the *Mki67* proliferation marker and metabolic genes, including *Glut4, Gck, Cd36*, and *Cpt1b* (**Fig.4I**). The tumors of mice implanted with *Ucp1*-CRISPRa adipose organoids had less Ki67+ cells than control mice (**Fig.4J**). These tumors also had lower CA9+ area per image, and CD31+ area per image view, indicating reduced hypoxia and angiogenesis (**Fig.4I-K**).

Additionally, mice implanted with *Ucp1*-CRISPRa adipose organoids had significantly lower plasma insulin levels than control mice (**fig.S4H**). Taken together, our results indicate that our CRISPRa-modulated adipose organoids could have systematic therapeutic effects in suppressing cancer growth.

### *UCP1-CRISPRa* upregulation of adipocytes from dissected breast tumors suppresses cancer growth

To further demonstrate the therapeutic potential of AMT, we treated adipocytes obtained from dissected human breast tissues with *UCP1*-CRIPSPRa AAV9 and tested their ability to suppress tumor progression by co-culturing them with breast cancer organoids generated from dissected breast tumors or metastatic pleural effusions (**Fig.5A**). Samples were obtained from patients who underwent breast surgery or thoracentesis. For the co-culture experiments, we used five different breast cancer organoids generated from patients with early-stage or metastatic triple-negative breast cancer [estrogen receptor (ER)-negative, progesterone receptor (PR)-negative/human epidermal growth factor receptor 2 (HER2)-negative] or hormone receptor-positive (ER-positive and/or PR-positive)/HER2-negative breast cancer (**Table S1**). Organoids from breast tumor tissue were generated by digesting cells for one hour in collagenase and organoids from metastatic pleural effusions were generated by isolating tumor spheroids via centrifugation, before in both cases embedding cancer cells in organoid culture using established protocols that enable long-term propagation of tumor organoids (*44*). In parallel, primary human adipocytes were isolated from human breast tissue using an established protocol (*45*). For the triple-negative breast cancer organoids derived from primary tumors, we used two different cases, including one where we generated organoids and isolated mammary gland adipose tissue from the same individual (TOR41). Adipocytes were infected with dCas9-VP64 only (negative control) or *UCP1*-CRISPRa AAV9. After five days, they showed strong *mCherry* expression, a fluorescent marker that is part of the gRNA AAV9 (**fig.S5A**), suggesting that they can be readily infected by our AAVs. In addition, gene analysis showed that both *mCherry* and *UCP1* expression levels were significantly higher in the *UCP1*-CRISPRa treated adipose organoids compared to the control organoids (**fig.S5B**).

**Fig. 5.**
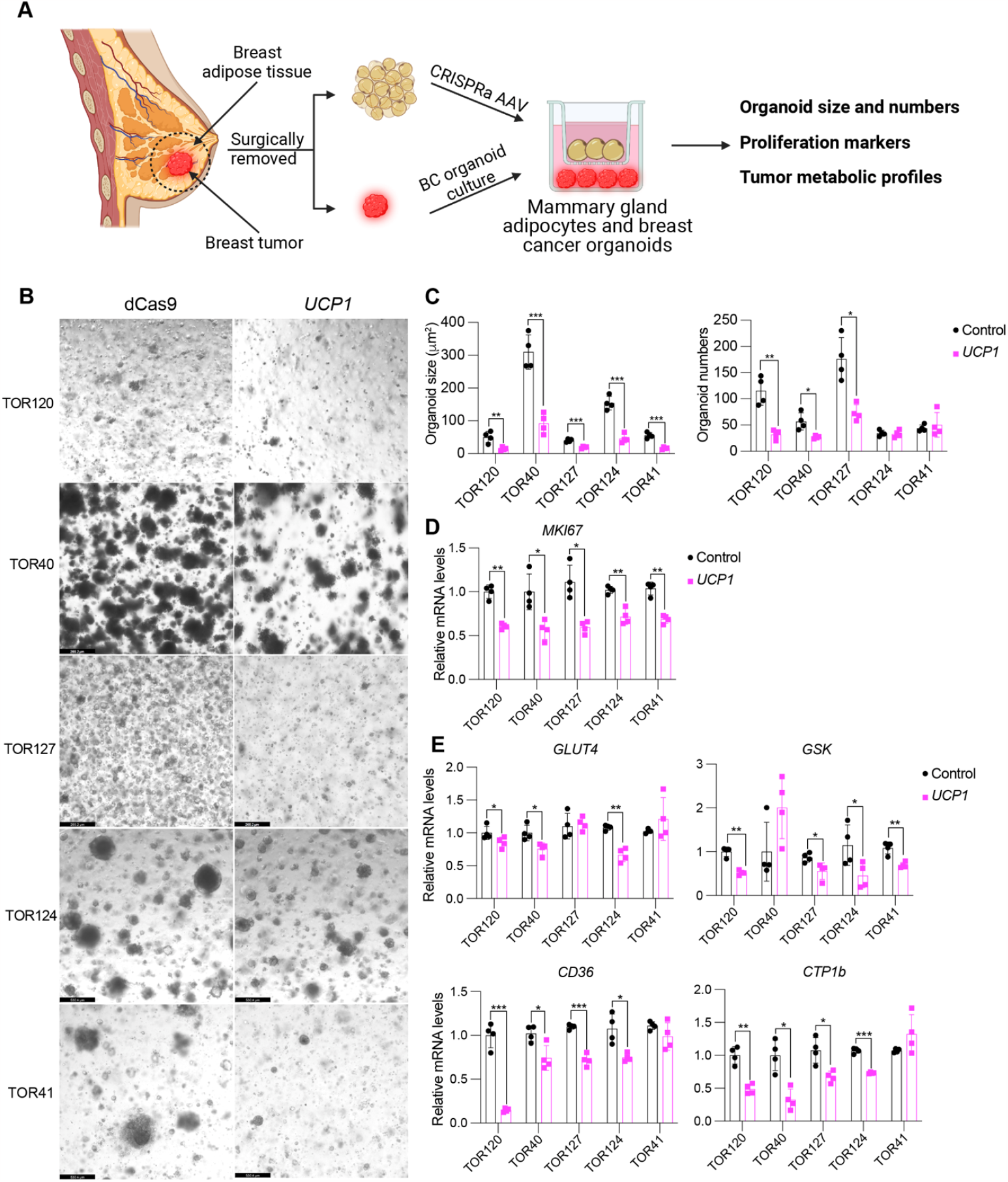
Cancer organoids co-cultured with *UCP1*-CRISPRa adipocytes, both from dissected breast tissue, lead to tumor suppression. (**A**) Schematic of the co-culturing model of *UCP1*-CRISPRa modulated human mammary adipocytes and breast cancer organoids from dissected breast tumors. (**B**) Representative images of breast tumor organoids from five dissected breast tumors that were co-cultured with *UCP1*-CRISPRa adipocytes or control (dCas9-VP64 only) adipocytes. (**C**) Breast cancer organoid size and numbers. Data are represented as mean ± S.D *≤0.05, **≤0.01, ***≤0.001. (**D**) qRT-PCR of proliferation marker gene, *MKI67*. Data are represented as mean ± S.D *≤0.05, **≤0.01. (**E**) qRT-PCR of metabolic genes, including *GLUT4, GCK, CD36*, and *CPT1b* of breast cancer organoids that were co-cultured with CRISPRa-modulated adipocytes. Data are represented as mean ± S.D *≤0.05, **≤0.01, ***≤0.001.

To test whether *UCP1*-CRISPRa AAV9 adipocytes can suppress cancer growth, we added them to a hydrogel dome on top of the breast cancer organoids with a fully-defined organoid medium (*44*) and co-cultured for five days (**Fig.5A**). In all five cases, we observed significant reduction in cancer organoid size (**Fig.5B**) and in most organoids also a reduction in organoid number (TOR124 and TOR41 were not significant) upon incubation with adipocytes infected with *UCP1*-CRISPRa AAV9 compared to the dCas9-VP64 only treated cells (**Fig.5C**). All five cancer organoid cases co-cultured with *UCP1*-CRISPRa had significantly lower proliferation marker *MKI67* expression compared to negative control (**Fig.5D**). We also observed significantly decreased levels of *GLUT4, GCK, CD36*, and *CTP1b* for the *UCP1*-CRISPRa co-cultured cancer organoids (**Fig.5E**), suggesting they have lower glycolysis and fatty acid metabolism. Overall, our results demonstrate that human adipocytes from a dissected breast tissue can upregulate *UCP1* via CRISPRa AAV9 and are able to reduce glycolysis and fatty acid metabolism and suppress breast cancer organoid growth. In addition, using matched mammary gland adipose tissue and breast cancer samples, we demonstrate the potential clinical utility of an *ex vivo* autologous transplantation of CRISPRa-modulated adipocytes to treat cancer.

## Discussion

Cancer cells are fast-proliferating cells that require large amounts of nutrients, including glucose and fatty acids. They can reprogram metabolic pathways to utilize available substrates in the surrounding environment. Targeting their metabolism can be a potent cancer treatment. Here, we developed CRISPRa-modulated human adipocytes that have increased glucose and fatty acid utilization by upregulating *UCP1, PPARGC1A*, and *PRDM16*. These CRISPRa-modulated adipocytes were able to suppress cancer growth in five different cancer cells lines, including two breast lines, colon, pancreas, and prostate. Co-transplanting CRISPRa-treated human adipose organoids and xenografts in mice led to a significant reduction in tumor growth in all five cancer types. These tumors had lower glycolysis and fatty acid metabolism and showed decreased hypoxia and angiogenesis. Implanting CRISPRa-modulated adipose organoids into genetic mouse models of breast and pancreatic cancers also prevented tumor development in mice. Furthermore, we demonstrated that implantation of CRISPRa-modulated adipose organoids distant from the cancer tissue leads to similar results. In addition, co-culturing cancer organoids from breast tumors and CRISPRa-modulated mammary adipocytes dissected from patients resulted in lower cancer growth. Combined, our data suggests that AMT has enormous potential to treat a wide variety of cancers.

Adipocytes offer a unique *ex vivo* therapeutic system with many of the needed procedures already established in the clinic. Liposuction and fat transplantation are commonly used in many surgical procedures, such as aesthetic and reconstructive surgery. Due to successful engraftment, adipose tissue transplantation has progressively evolved, not only in plastic and reconstructive surgery, but also for therapeutic treatments (*46*). Several reports using rodent models have shown that BAT transplantation has beneficial metabolic outcomes (*47-50*). These also include the use of *UCP1-*CRISPRa modulation in human white preadipocytes to induce browning, followed by their transplantation in mice treated on a high-fat diet, leading to improved body weight, glucose tolerance and insulin sensitivity (*34*). Our work further showcases how these ‘brown’ adipocytes and adipose organoids can be utilized for cancer treatment.

There is a growing interest in carrying out human adipose tissue grafting by using adipose stem cells or progenitors due to their resistance to trauma and long-term survival following transplantation. One such example, is the use of CRISPRa to upregulate the relaxin family peptide receptor 1 (*RXFP1*) in adipose-derived stem cells and their transplantation in a diabetes mellitus induced erectile dysfunction rat model, showing amelioration of the erectile disfunction phenotype (*51*). Organoids could be advantageous for these cells as they provide a 3D-culture that can better recapitulate the heterogeneity of adipose tissue, respond better to endogenous stimuli and form tissue microenvironments that could more efficiently integrate with cancer cells following transplantation. Several groups have successfully grown adipose organoids from mouse adipose stem cells (*33, 40, 52, 53*). Here, we were able to culture human adipose organoids from preadipocytes using various conditions from previous studies (*33, 40, 52, 53*). These human adipose organoids exhibited mature adipocytes markers, including *FABP4* and *PLIN1*. In addition, in our hands they could last up to a year after being subcutaneously implanted in mice, showing the potential for long-term treatment (data not shown). Finally, in our study we show that their CRISPRa modulation could significantly reduce tumor size, glycolysis and fatty acid metabolism and improve hypoxia and angiogenesis in cancer mouse models. Further development of these organoids and their modulation could likely improve these attributes and their therapeutic use for a wide range of diseases.

Our work builds on the recent observation that activating BAT, via cold exposure, increases adipocyte glucose uptake, lipid metabolism and significantly inhibits tumor progression (*15*). Here, instead of placing tumor models in cold conditions, we took advantage of CRISPRa to increase the expression of BAT key regulators, including *UCP1, PPARGC1A, PRDM16* to engineer adipocytes to have increased glucose and fatty acid uptake and metabolism, and used them to deplete resources from cancer cells. Interestingly, and in line with the BAT cold activation results (*15*), implanting CRISPRa-modulated adipose organoids distal from the tumors also led to suppression of cancer growth, suggesting that complex surgical procedures for tumors with limited access might not be needed for this approach. This phenomenon might be due to additional mechanisms in which CRISPRa-modulated adipose organoids can suppress cancer growth. Previous studies have shown that hyperinsulinemia can lead to cancer growth due to insulin being a powerful mitogen and survival factor (*54-56*). For example, administration of Dapagliflozin, a SGTL2 inhibitor that lowers blood glucose, and a controlled-release mitochondrial protonophore (CRMP) suppress cancer growth in mice by reversing hyperinsulinemia (*57*). Since BAT is widely known to reduce whole-body blood glucose and insulin levels in humans (*15, 49, 58*), we hypothesize that the CRISPRa-modulated adipose organoids can reduce cancer progression by lowering plasma insulin levels. Indeed, our data, using both xenograft and genetic mouse models, shows that mice implanted with CRISPRa-modulated adipose organoids have reduced plasma insulin levels compared to dCas9-V64 control mice (**fig.S3G, S4F**,**H**).

Amongst BAT-activating genes, *UCP1* showed the most robust effect in terms of cancer suppression. It would be interesting to further develop this AMT approach to upregulate additional genes that could aid in cancer therapy. These could include for example upregulation of *GLUT1* and *GLUT4*, that are the main glucose transporters in adipose cells, with *GLUT4* being the most abundant and insulin responsive (*59*). Glucose metabolism associated genes, such as the transcription factor *FOXO1* (*60*) and the G-coupled receptors, GPR20 and GPR120, which were implicated in improved glucose uptake and insulin resistance (*61, 62*), and *AIFM2*, which promotes glycolysis in BAT (*63*). Fatty acid oxidation associated genes, including for example the fatty acid transporter *CD36*, a key transporter for fatty acid oxidation, *CPT1b*, and the fatty acid breakdown enzyme, *ACC1*. Additional modifications could also be engineered in these adipocytes/adipose organoids, including for example utilization of their endocrine capabilities to secrete chemotherapeutic drugs or other cancer therapeutic associated compounds.

In this study, we used AAV-based CRISPRa to upregulate genes. However, it is worth noting that both upregulation and delivery could also be carried out using other modalities. For example, gene upregulation could be carried out using zinc fingers, TALENS, generation of specific mutations via regular CRISPR editing or base or prime editing in promoters or enhancers or standard overexpression using a cDNA mammalian expression construct of the gene of interest. Delivery could also be carried out with other viruses, such as lentivirus that is widely used for CAR-T therapy, but has a major caveat of genomic integration, or various non-viral nucleic acid delivery vehicles such as nanoparticles (*64*) or virus-like particles (VLPs) (*65*). Various drugs could also be used to upregulate specific genes in adipocytes in a global manner in cancer patients. In addition, downregulation of certain genes in adipocytes or adipose organoids, using CRISPRi, siRNA, CRISPR editing or other techniques could also be utilized for AMT. For example, a recent study used CRISPR to deplete the nuclear receptor interacting protein 1 (*NRIP1*) to make ‘brown’ adipocytes which upon implantation in mice decreased their adiposity on a high-fat diet (*66*).

A link between obesity, excess amount of WAT and cancer development and progression has been established, with nearly 40% of all cancer deaths in the United States being attributed to obesity (*67*). There are numerous proposed mechanisms in how WAT is linked to cancer development and progression, including chronic inflammation, hyperinsulinemia, steroid hormones, and adipokines (*68-74*). In addition, in glucose-rich conditions, cancer cells synthesize *de novo* FAs from intermediates of the glycolysis-TCA cycle (lipogenesis)(*75, 76*). The synthesized FAs are then utilized to synthesize triglycerides and are stored as lipid droplets in cancer cells. When energy is needed, these lipid droplets undergo lipolysis to release FAs which are subjected for b-oxidation (*76-78*). Since BAT is highly associated with improved glucose tolerance and insulin sensitivity (*79*), one could envision a personalized treatment development for obese cancer patients, whereby AMT is used not only to target cancer and its unique metabolism in these patients but also treat their metabolic disease. One major hurdle in our approach that needs to be taken into account is cancer-associated cachexia (*41*). While we did not observe weight loss in our mouse models, a longer treatment time could possibly lead to a reduction in body weight, as was shown for *UCP1*-CRISPRa mice on a high-fat diet (*34*). Modification of other genes than *UCP1*, removal of the implant after a certain time or having a molecular kill switch could all be potential solutions for cachexia if observed.

In summary, our results provide proof-of-principle results for a novel cancer therapeutic approach, termed AMT, that can be further improved and personalized for specific cancers and patients. Similar to chimeric antigen receptor (CAR)-T cell therapy, AMT can be readily used in the clinic as cells can be obtained from cancer patients via liposuction or other procedures, engineered and transplanted back into the same individual for therapeutic benefit. The utilization of adipocytes from dissected breast tissue, as performed in our study, further showcases the clinical utility of such an *ex vivo* approach. In particular, for breast cancer this could be particularly straightforward as many mastectomies are followed up by reconstructive surgery with autologous tissue (*80*), which could be manipulated priorto this procedure. Unlike T-cells, adipocytes have a lower immune response (*34, 81*) which could allow more straightforward development of ‘off-the-shelf’ adipocytes or adipose organoids for cancer and other treatments. Their ease of growth in culture, long lasting and robustness, and lower multiplicity (*82*) along with existing clinical procedures to remove and transplant them make them an exceptional cell type for cancer and other cellular-based disease therapies.

## Acknowledgments

We would like to thank Dr. Hei Sook Sul for kindly providing the human preadipocyte cell line and the patients that provided clinical material. We would also like to thank Drs. Felix Feng and Alan Ashworth at UCSF for their support and advice. The content is solely the responsibility of the authors and does not necessarily represent the official views of the National Institutes of Health.

## Author contributions

Conceptualization: HPN, NA

Methodology: HPN, RS, MB, HH, EM, K.A

Samples: JMR, FL, DD, MM, LH, LE

Writing, Review & Editing: HPN, NA with contributions from all authors

Visualization: HPN, NA

Supervision: NA

Funding acquisition: NA, JMR

## Data and materials availability

All data are available upon request. Requests for materials should be directed to NA.

