## Supplemental material for "Implantation of engineered adipocytes that outcompete tumors for resources suppresses cancer progression"

Materials and Methods

Human and mouse adipocytes

For adipocyte differentiation of human preadipocytes, human preadipocytes or mouse 3T3-L1 preadipocytes (both kind gifts from Dr. Hei Sook Sul, UC Berkeley) were cultured to 100% confluency in DMEM, supplemented with 10% FBS and fresh media were replaced. After 48 hours, cells were subjected to adipocyte differentiation by adding a cocktail of 3-isobutyl-1-methylxanthine (IBMX) (0.5M) (Sigma-Aldrich; 410957), dexamethasone (1uM) (Sigma-Aldrich; D1756), and insulin (10ug/ml) (Sigma-Aldrich; I9278). Media was replaced every two days with insulin containing DMEM complete media (Fisher Scientific; 11-965-118) during differentiation.

**Cancer cell lines**

All the cancer cells were acquired from American Type Culture Collection (ATCC). MCF7 (ATCC; HTB-22) was cultured in Eagle’s minimum essential medium (ATCC; 30-2003), supplemented with 10% FBS and 10ug/ml human recombinant insulin (Sigma-Aldrich; I9278). MDB-MA-436 (ATCC; HTB-130) was cultured in Leibovitz’s L-15 medium (ATCC; 30-2008) with 10ug/ml insulin,16 ug/ml glutathione (Sigma-Aldrich; G6013) and 10% FBS. SW-1417 (ATCC; CCL-238) was grown in Leibovitz’s L-15 medium supplemented with 10% FBS. Panc 10.05 (ATCC; CRL-2547) was cultured in RPMI-1640 medium (ATCC; 30-2001) with 10ug/ml human insulin and 15% FBS. DU-145 (ATCC; HTB-81) was cultured in Eagle’s minimum essential medium with 10% FBS.

**CRISPRa AAV *in vitro* optimization**

Five gRNAs targeting the promoter of human *UCP1*, *PPARGC1A*, and *PRDM16* or mouse *Ucp1* were designed using the Broad Institute CRISPick Tool (*83*) (**Table S2**). These guides were individually cloned into pAAV-U6-sasgRNA-CMV-mCherry-WPREpA (*84*) at the *BstXI* and *XhoI* restriction enzyme sites using the In-Fusion (Takara Bio; 638910) cloning methods as described in (*84*). 5x10^5^ human preadipocytes or mouse 3T3-L1 cells were plated into 12-well plate and were then subjected to adipocyte differentiation protocol after two days of confluency. At day four of differentiation, cells were transfected with 0.5ug of dCas9-VP64 and 0.5ug of gRNA plasmid using X-tremeGene (Sigma; 6366236001) and CombiMag reagent (OZ Biosciences; CM21000) following the manufacturer’s protocol. At day eight of differentiation, cells were lysed with Trizol and RNA was collected and cDNA and qRT-PCR were performed, as mentioned in RNA Isolation and qRT-PCR section. The two gRNAs with the highest upregulation for each gene were packaged into rAAV-9 serotype virons. rAAV-9 serotype virons were produced by transfecting AAVpro 293T cell (Takara; 6322723) with pCMV-sadCas9-VP64 (Addgene; 115790) or pAAV-U6-sasgRNA-CMV-mCherry-WPREpA (*84*) along with packaging vectors, including PAAV2/9n (Addgene; 112865) and pHelper vectors using TransIT293 reagent (Mirus; 2700). After 72 hours, AAV particles were collected and purified using AAVpro Cell & Sup. Purification Kit Maxi (Takara; 6676) and quantified by the AAVpro Titration Kit (Takara; 6233). gRNA AAV viruses (1x10^6^ MOI) and dCas9-VP64 AAV (1x10^6^ MOI) were used to infect human and mouse differentiated adipocytes. After five days, RNA was collected, cDNA was made, and qRT-PCR was done as described below.

**RNA Isolation and qRT-PCR**

Total RNA from sorted or cultured cells was extracted using Trizol reagent (ThermoFisher; 15596026). Reverse transcription was performed with 1μg of total RNA using qScript cDNA Synthesis Kit (Quantabio; 95047) following the manufacturer’s protocol. qRT-PCR was performed on QuantStudio 6 Real Time PCR system (ThermoFisher) using Sso Fast (Bio-Rad; 1705205). Statistical analysis was performed using ddct method with *GAPDH* or *Gapdh* primers as control (see primer sequences in **Table S2**). Gene expression results were generated using mean values for over n=4-6 biological replicates.

**Seahorse assay**

Human preadipocytes (5x10^5^ cell/well) were plated in 12-well tissue culture plates and subjected to adipocyte differentiation protocol. Adipocytes and cancer cell lines were trypsinized and reseeded in XF96 plates (Agilent; 102905-100) at 20K cells per well and assayed the next day. On the day of experiments, the cells were washed two times and maintained in XF base medium (Agilent; 1033334) supplemented either with 1 mM sodium pyruvate (ThermoFisher; 11360070) and 17.5 mM glucose (Sigma-Aldrich; G7021), for the mitochondria stress test or with 2 mM glutamine (StemCell technologies; 07100) for the glycolysis test. For FAO tests, cells were washed and incubated with substrate-limiting medium DMEM (Corning; 17-207-CV) supplemented with 0.5 mM glucose, 1 mM glutaMAX (ThermoFisher; 3050061), 0.5 mM carnitine (Sigma-Aldrich; C0283), and 1%FBS. Cells were then incubated at 37^o^C in a non-CO2 incubator for 1 hour. The assay was performed using Seahorse XFe96 analyzer (Agilent; XFe96). Oxygen consumption measured under 1.5 μM oligomycin (Sigma-Aldrich; 75351), 2 μM FCCP (Sigma-Aldrich; C2920), and 0.5 μM of Rotenone (Sigma-Aldrich; R8875)/Antimycin (Sigma-Aldrich; A8674). Extracellular acidification rate (ECAR) was measured under 10 mM glucose, 1 uM oligomycin, and 50 mM 2-deoxyglucose (Sigma-Aldrich; D8375).

**Glucose uptake assay**

Human preadipocytes (3x10^5^ cell/well) were plated in 96-well tissue culture plates and subjected to adipocyte differentiation protocol. One day before the assay, media was replaced with serum-free media. On the day of the assay, using glucose uptake-Glo assay (Promega; J1342), cells were incubated with 1 nM insulin (Sigma-Aldrich; I9278) for 1 hour prior to the assay following the manufacturer's protocol.

**Co-culturing CRISPRa-modulated adipocytes with cancer cells**

5x10^5^ human preadipocytes were plated in 12-well tissue culture plates and subjected to adipocyte differentiation protocol. At day 2 of differentiation, cells were transduced with gRNA AAV viruses (1x10^6^ MOI) and dCas9-VP64 AAV (1x10^6^ MOI). After 6 days, cells were collected and replated into the upper-well of 12-well transwell plate (Fisher Scientific; 07-200-150) in which 3x10^5^ cancer cells were plated in the lower-well a day earlier. The cells were culture in adipocyte differentiation media, described previously, and designated media for each cancer cell line (1:1 ratio). After three days, cancer cells were collected for imaging and seahorse assay. RNA was collected, and cDNA was made, and qRT-PCR was carried out as previously described. Differential expression was determined using the ddct method with *GAPDH* primers as control (primer sequences in **Table S2**).

**Human and mouse adipose organoid culture**

Human or mouse 3T3-L1 preadipocytes (0.5x10^6^ cells) were plated in 96-well Nunclon Sphera ULA U-bottom plates in (ThermoFisher; 174929). Organoids formed after 48 hours and were then differentiated to adipose organoids using a differentiation cocktail containing 3-isobutyl-1-methylxanthine (IBMX) (0.5M), dexamethasone (1uM), and insulin (10ug/ml). Adipocytes formed 21-days post differentiation. Human adipose organoids were then transduced with gRNA AAV viruses (1x10^6^ MOI) and dCas9-VP64 AAV (1x10^6^ MOI). After 5 days, organoids were collected and mixed with Matrigel (Corning; 354234) and subcutaneously injected into mice (10 organoids/mouse).

**Co-transplantation of CRISPRa-modulated human adipose organoids and cancer cells**

All animal studies were carried out in accordance with the University of California San Francisco Institutional Animal Care and Use Committee protocol number AN197608. Mice were housed in a 12:12 light-dark cycle, and chow diet (Envigo; 2018S) and water were provided *ad libitum*. Five-week-old immune-deficient SCID mice (JAX; 001303) were anesthetized using isoflurane and subcutaneously injected with 2x10^6^-6x10^6^ cancer cells in phosphate buffered saline (PSB). After 6-12 weeks, depending on the cancer cell lines, we subcutaneously injected CRISPRa AAV human adipocytes or adipose organoids to a site adjacent to the tumor. Following 4-6 weeks, mice were euthanized, and tumors and adipose implants were collected. We measured tumor size with calipers and tumor volume according to standard formula (length x width^2^ x 0.52).

**Immunofluorescence**

A portion of isolated tumors were fixed with 1% paraformaldehyde for two hours and prepared for cryostat sectioning. Each sample were cut into 5 umm thick sections and blocked with 3% BSA (Miltenyi Biotech; 130-091-376) blocking solution for 30 minutes. For Ki-67 staining, slides were permeabilized with 1% triton X-100 PBS solution for 15 minutes and washed with 0.1% Tween20 PBS solution prior to blocking. Slides were then incubated with primary antibodies (information and concentration listed in **Table S3**) overnight at 4^0^C and washed 3x with 0.1% Tween20 PBS. The slides were then incubated with secondary antibodies (information and concentration listed in **Table S3**) for 60 minutes at room temperature and then washed 3x with 0.1% Tween20 PBS. Images were obtained using a confocal microscope (Zeiss; LSM880) and analyzed with Fiji (*85*).

**CRISPRa-modulated mouse adipose organoid implantation in pancreatic and breast cancer genetic mouse models**

KPC, tamoxifen-inducible" is a **K**ras^LSL-G12D^;**p**53^LoxP^;Pdx1-**C**reER triple mutant model of tamoxifen-inducible pancreatic ductal adenocarcinoma (PDAC) on an C57/BL6 background was acquired at the Jackson Laboratory (032429). To induce *Cre* recombination, tamoxifen was administered to pups via lactation following oral gavage of the mother with 6 mg tamoxifen (suspended in corn oil) at postnatal days 0, 1, 2 and 4. We orthotopically implanted mouse adipose organoids (10 organoids/mouse) mixed with Matrigel (Corning; 354234) near the pancreas in 4-week-old mice, using a similar protocol describe in (*86*) and dissected the pancreas 6 weeks after the implantation. The *MMTV-PyMT* breast cancer mice were acquired from the Jackson Laboratory (002374). Four-week-old female mice were implanted with mouse adipose organoids (10 organoids/mouse) mixed with Matrigel (Corning; 354234) and subcutaneously injected into mice either at the third nipple or the back. The tumors were collected six weeks post implantation.

**Cancer organoids**

Organoids were generated from breast tumor tissue or tumors from malignant effusions. Patients were consented for specimen collection using IRB-approved tissue collection protocols at the University of California, San Francisco and the Brigham & Women’s Hospital. Surgical specimens were obtained from the UCSF Medical Center or Brigham & Women’s Hospital on the day of their procedure, viably frozen as tissue pieces, or used to generate formalin-fixed, paraffin-embedded (FFPE) sections or organoid cultures. Each tissue was minced using razor blades and digested in a solution containing DMEM/F12 (Gibco; 11330), 2 mM GlutaMax (Gibco; 35050), 10 mM HEPES (Gibco; 15630), 50 U/mL Penicillin-Streptomycin (Gibco; 15070), and 1 mg/ml collagenase XI (Sigma-Aldrich; C9407). Tissue digestion was performed at 37°C with constant shaking at 150 rpm for 1-2 hours. Cells were then pelleted by centrifugation, further dissociated by sequentially pipetting with 10, 5, and 1-ml pipette tips, and recentrifuged. The resulting cell pellet was used directly to establish organoid cultures by embedding in basement membrane extract, allowing this to harden at 37°C for 20 minutes to form a hydrogel dome, and then overlaying this dome with Type 1 Organoid Medium as previously describe (*87, 88*). All metastatic cancer organoids were derived from malignant pleural effusions that were collected from metastatic breast cancer patients undergoing thoracentesis at the UCSF Medical Center. The fluid samples were placed on ice within four hours after collection. Organoids were generated by washing malignant effusions with PBS, collecting tumor spheroids by centrifugation, and incubating with 3-5 ml RBC lysis buffer (BioLegend; 420301) for 10-15 minutes when there were visible RBCs, followed by embedding the cell pellet in organoid culture as described above.

**Mammary gland adipocyte isolation**

For adipocyte extraction, mammary gland adipose tissues from excess tissue removed during breast surgeries were washed with PBS three times and mechanically minced into smaller pieces. The tissue mixture was incubated with PBS with 3% BSA and Collagenase I (1mg/ml) for 45 minutes at 37^o^C, centrifuged and the top lipid layer was discarded. The remaining mixture was filtered twice through a 200um strainer and centrifuged. Mature adipocytes were isolated and washed twice with PBS and were plated in suspension in 6 well plates in high glucose DMEM media supplemented with 10% FBS. For transduction of CRISPRa, *UCP1* AAV9 viruses (1x10^6^ MOI) and dCas9-VP64 AAV9 (1x10^6^ MOI) were mixed with 20ul of AdeoMag (OzBiosciences, AM71000) and adipocytes and incubated for 15 minutes. The virus and cell mixture were distributed onto 6-well plate placed upon the magnetic plate and incubated for 15 more minutes.

**Co-culturing of cancer organoids and adipocytes**

Five days post infection of CRISPRa AAV9, adipocytes were placed in a tissue transwell culture plate (Fisher Scientific; 07-200-150) of tumor organoids in 1:1 ratio of adipocyte media and breast cancer organoid media. Cells were incubated for up to seven days and then examined for adipocyte and tumor phenotypes.

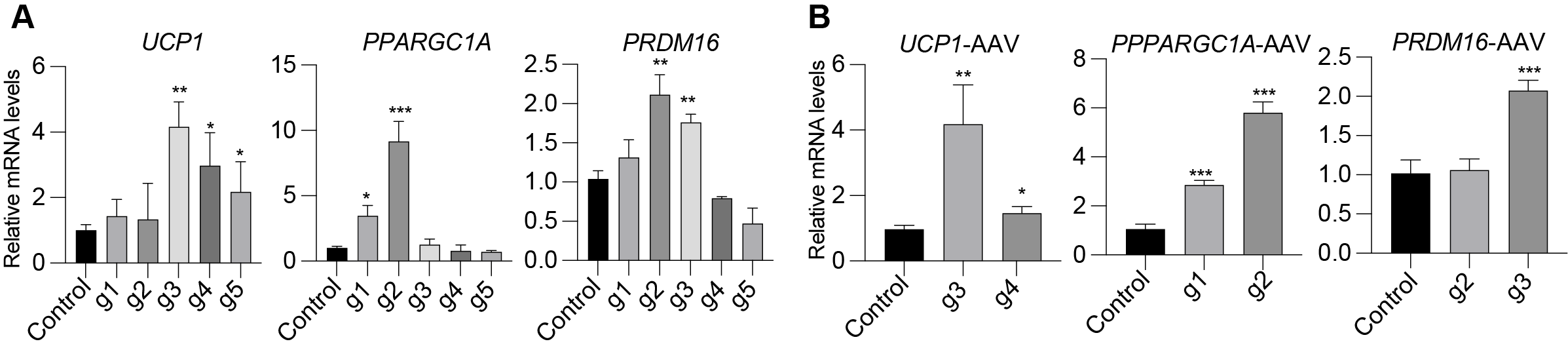

**Fig. S1. CRISPRa upregulation of *UCP1*, *PPARGC1A*, and *PRDM16* in human white adipocytes.** (**A**) qRT-PCR of *UCP1*, *PPARGC1A*, and *PRDM16* in white adipocytes transfected with CRISPRa using five different gRNAs per gene. Data are represented as mean ± S.D. *≤0.05, **≤0.01, ***≤0.001. (**B**) qRT-PCR of *UCP1*, *PPARGC1A*, and *PRDM16* in human adipocytes transduced by AAV9 CRISPRa with the top two gRNAs per gene. Data are represented as mean ± S.D. *≤0.05, **≤0.01, ***≤0.001.

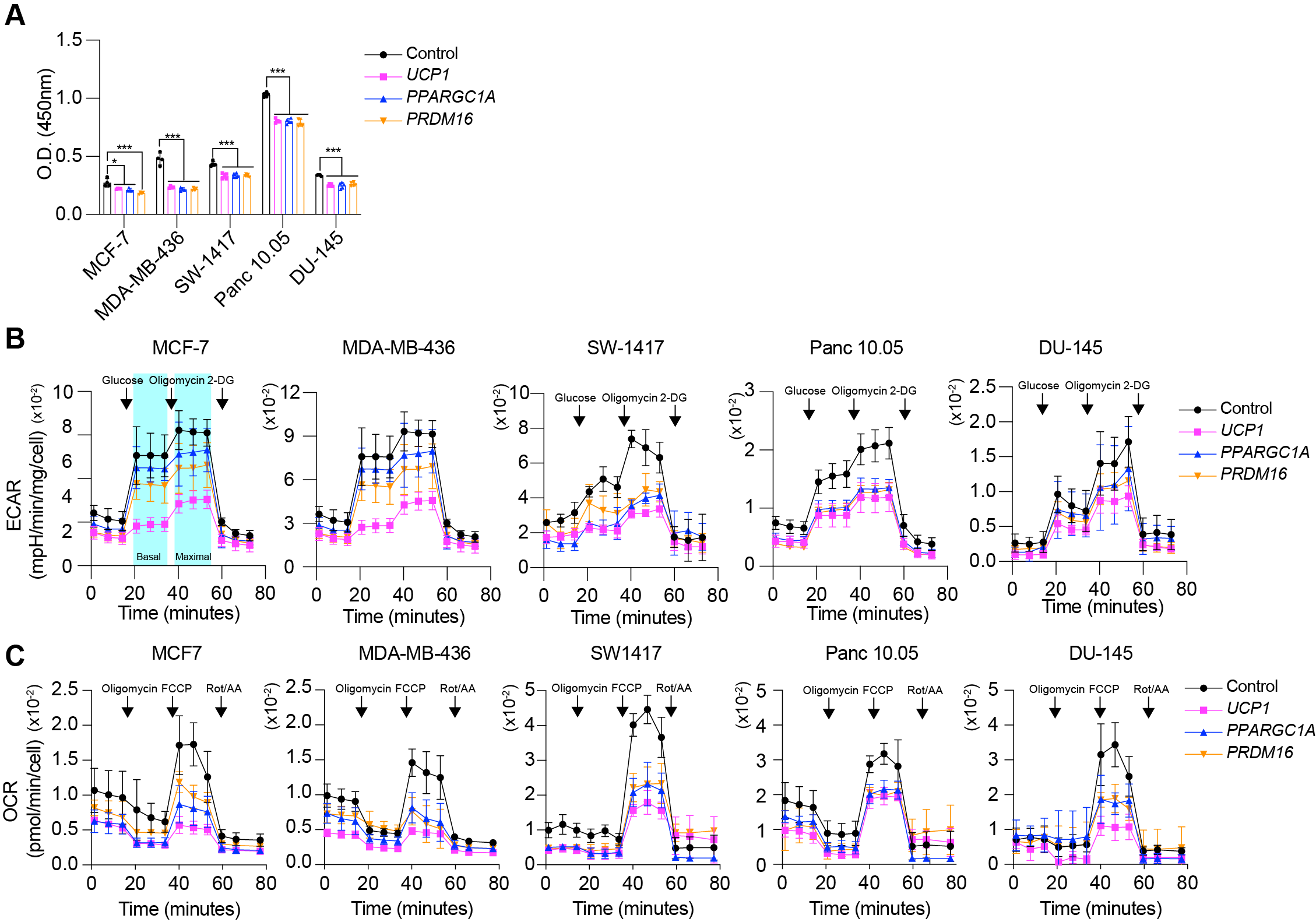

**Fig. S2. Proliferation and metabolic rate of various cancer cells co-cultured with CRISPRa-modulated human adipocytes.** (**A**) BrdU incorporation, (**B**) Oxygen consumption rate (OCR) ,and (**C**) Fatty acid oxidation of various cancer cells (MCF-7, MDA-MB-436, SW-1417, Panc.10.05, and Du-145) that were co-culture with CRISPRa-modulated adipocytes. Data are represented as mean ± S.D. *≤0.05, **≤0.01, ***≤0.001.

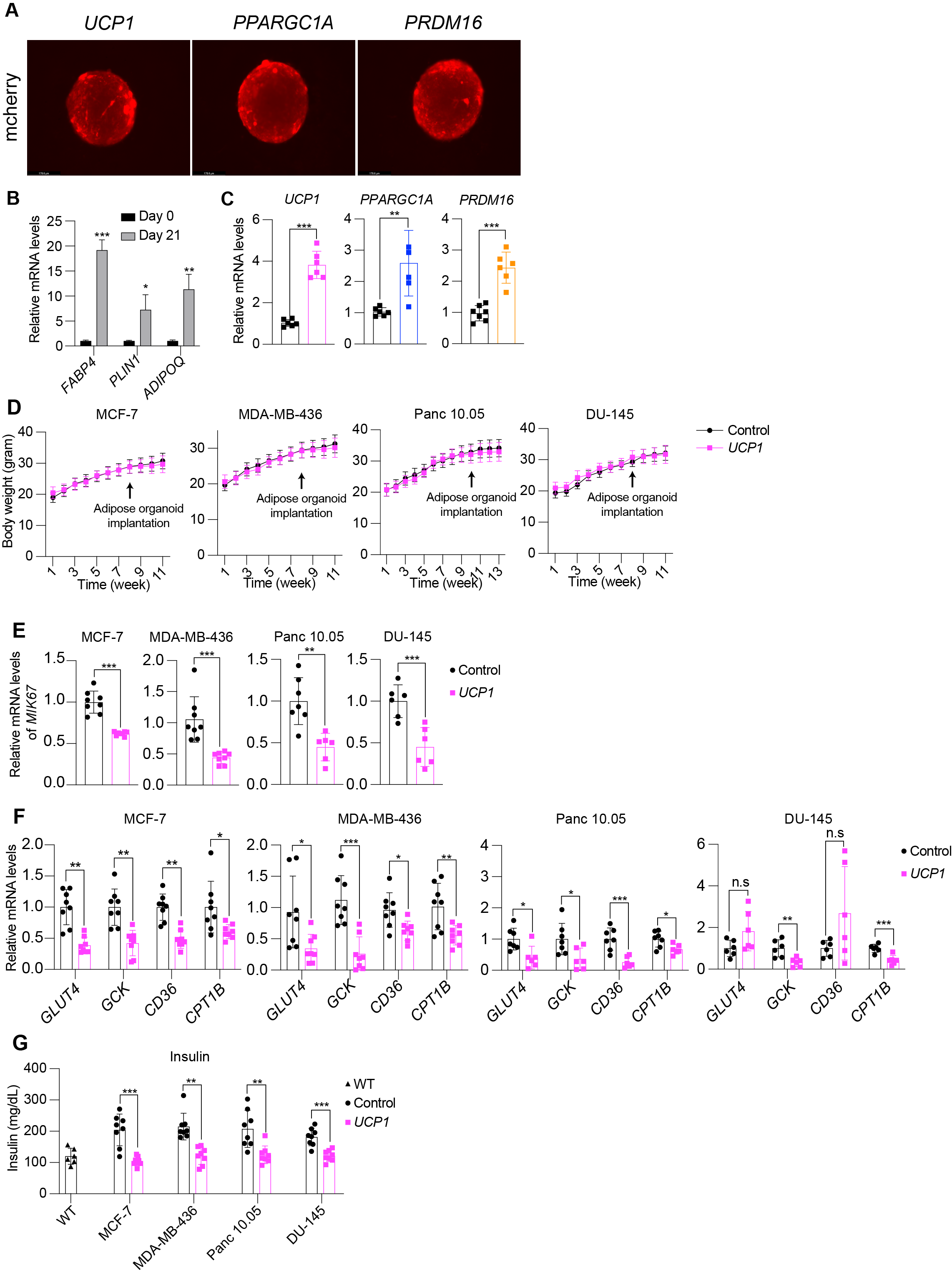

**Fig.S3. CRISPRa upregulation of *UCP1*, *PPARGC1A*, and *PRDM16* in human adipose organoids and proliferation and metabolic gene expression in tumors co-transplanted with CRISPRa-modulated adipose organoids.** (**A**) Representative images of human adipose organoids transduced with *UCP1*, *PPARGC1A*, and *PRDM16* AAV9 CRISPRa showing mCherry expression. (**B**) qRT-PCR of *FABP4*, *PLIN1*, and *ADIPOQ* in adipose organoids. Data are represented as mean ± S.D. *≤0.05, **≤0.01, ***≤0.001. (**C**) qRT-PCR of *UCP1*, *PPARGC1A*, and *PRDM16* in human adipose organoids transduced by AAV9 CRISPRa. Data are represented as mean ± S.D. **≤0.01, ***≤0.001. (**D**) Body weight of mice that were co-transplanted with *UCP1*-modulated adipose organoids and cancer cells. (**E**) qRT-PCR of *MKI67* in xenograft tumors derived from MCF-7, MDA-MB-436, Panc 10.05, and DU-145 cancer cells that were co-transplanted with *UCP1*-CRISPRa modulated adipose organoids. Data are represented as mean ± S.D. **≤0.01, ***≤0.001. (**F**) qRT-PCR of *GLUT4, GCK, CD36,* and *CTP1B* in xenograft tumors. Data are represented as mean ± S.D. *≤0.05, **≤0.01, ***≤0.001. (**G**) Plasma insulin levels of wild type (WT) SCID mice and mice that were co-transplanted with *UCP1*-CRISPRa adipose organoids and cancer cells. Data are represented as mean ± S.D. **≤0.01, ***≤0.001.

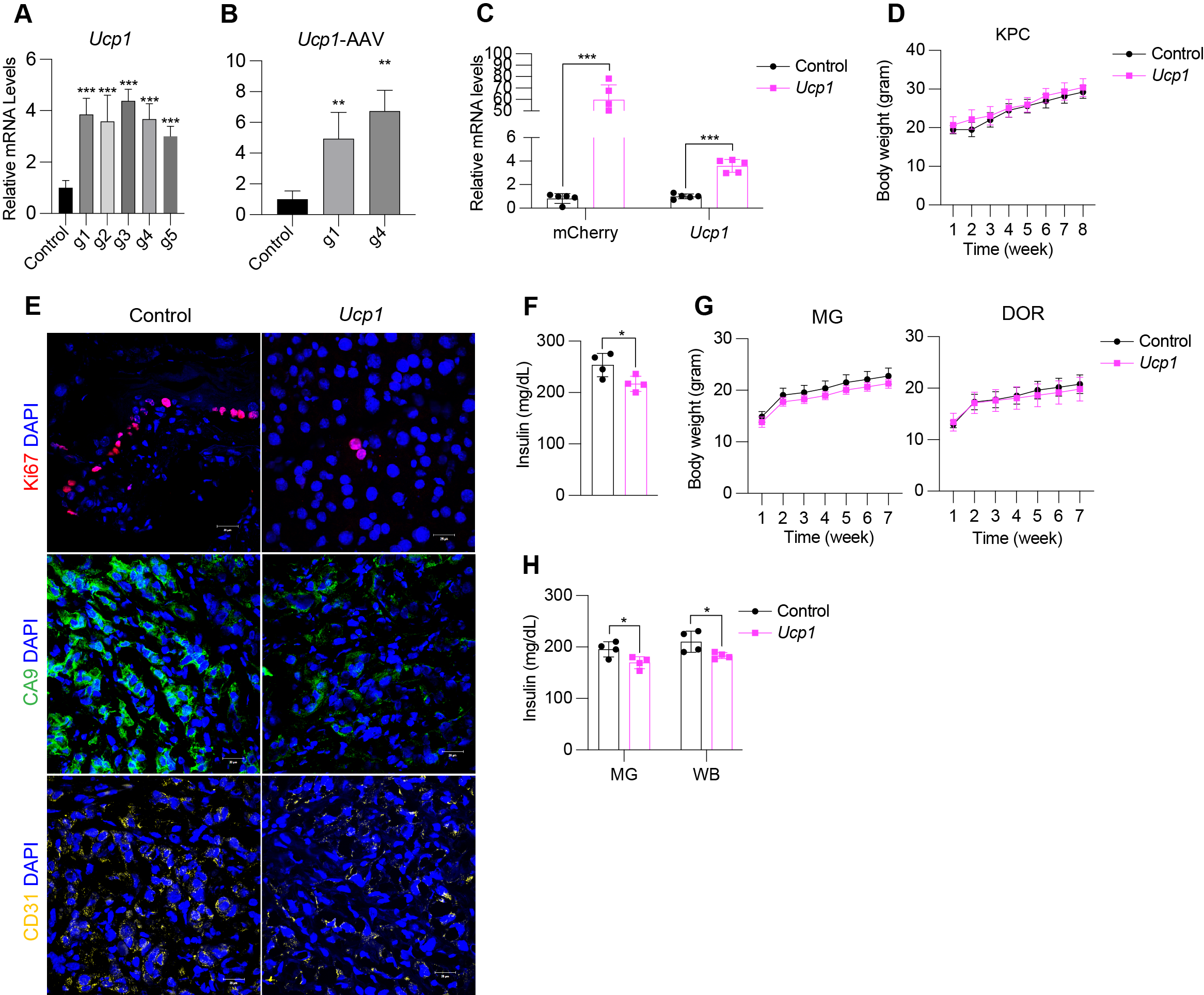

**Fig. S4. CRISPRa upregulation of *Ucp1* in mouse white adipocytes and phenotypic analysis of mouse genetic cancer models.** (**A**) qRT-PCR of *Ucp1*, in white adipocytes transfected with CRISPRa using five different gRNAs. Data are represented as mean ± S.D. *≤0.05, ***≤0.001. (**B**) qRT-PCR of *Ucp1* in mouse adipocytes transduced by AAV9-CRISPRa with the top two gRNAs per gene. Data are represented as mean ± S.D. **≤0.01. (**C**) qRT-PCR of mcherry and *Ucp1* in mouse adipose organoids. Data are represented as mean ± S.D. ***≤0.001. (**D**) Body weight of pancreatic cancer KPC mice implanted with control (dCas9-VP64) or *Ucp1*-upregulated mouse adipose organoids. (**E**) Immunofluorescence staining and quantification of Ki67, CA9, and CD31 in cryosections of pancreatic tumors. (**F**) Plasma insulin levels of pancreatic cancer genetic mice implanted with either *Ucp1*-CRISPRa adipose organoids or control organoids. Data are represented as mean ± S.D. *≤0.05. (**G**) Body weight of breast cancer (*MMTV-PyMT*) mice implanted with control (dCas9-VP64) or *Ucp1*-CRISPRa mouse adipose organoids nearby mammary gland (MG) or distal (DOR). (**H**) Plasma insulin levels of breast cancer genetic mice implanted with either *Ucp1*-CRISPRa adipose organoids or control organoids nearby mammary gland (MG) or distal (DOR). Data are represented as mean ± S.D. *≤0.05.

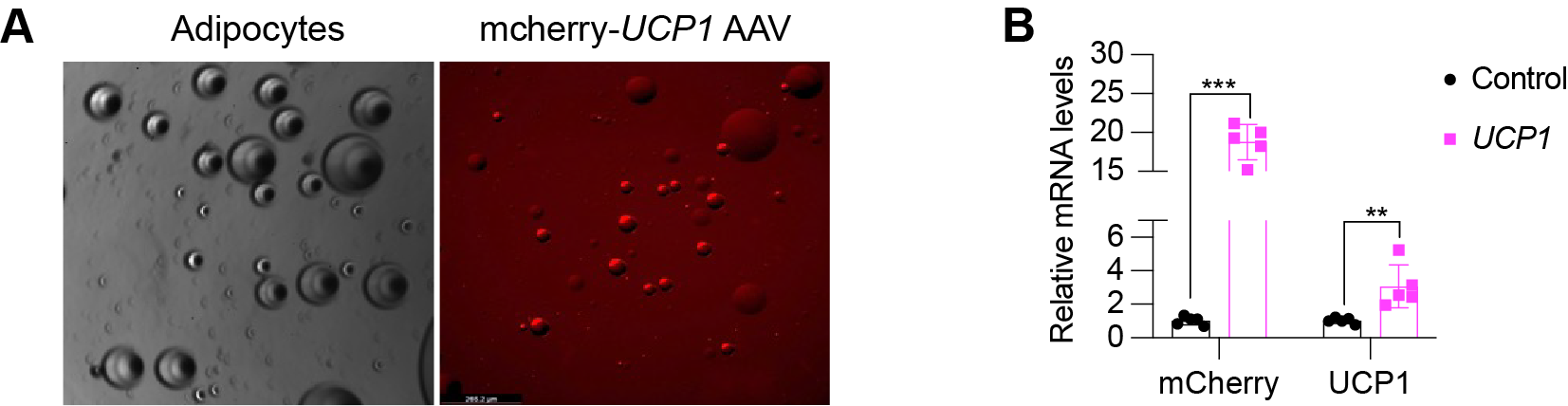

**Fig. S5. CRISPRa upregulation of *UCP1* in human mammary gland adipocytes.** (**A**) Representative images of adipocytes isolated from mammary glands that were transduced with *UCP1*-CRISPRa AAV9. (**B**) qRT-PCR of *dCas9, mcherry,* and *UCP1* in CRISPRa-modulated mammary gland adipocytes. Data are represented as mean ± S.D. *≤0.05, **≤0.01, ***≤0.001.

**Table S1.** Breast cancer organoid status.

| **Name** | **ER** | **PR** | **HER2** | **Metastatic** | **Inflammatory?** | **Source of adipocytes?** |
| --- | --- | --- | --- | --- | --- | --- |
| **TOR40** | Negative | Negative | Negative | no | yes | ad-1 (unmatched breast) |
| **TOR120** | Negative | Negative | Negative | yes | no | ad-2 (unmatched breast) |
| **TOR127** | Positive | Positive | Negative | yes | no | ad-3 (unmatched breast) |
| **TOR124** | Negative | Negative | Negative | yes | no | ad-4 (unmatched breast) |
| **TOR41** | Negative | Negative | Negative | no | yes | ad-5 (matched breast) |

**Table S2.** Sequences of primers for RT-qPCR, cloning, genotyping and gRNAs.

| **Gene/Use/sgRNA** | **Primers** |
| --- | --- |
| *GAPDH* | GTC TCC TCT GAC TTC AAC AGC G |
|  | ACC ACC CTG TTG CTG TAG CCA A |
| *UCP1* | TGG AAT AGC GGC GTG CTT G |
|  | CTC ATC AGA TTG GGA GTA G |
| *PPARGC1A* | CCA AAG GAT GCG CTC TCG TTC A |
|  | CGG TGT CTG TAG TGG CTT GAC T |
| *ADIPOQ* | CAG GCC GTG ATG GCA GAG ATG |
|  | GGT TTC ACC GAT GTC TCC CTT AG |
| *PLIN1* | GCG GAA TTT GCT GCC AAC ACT C |
|  | AGA CTT CTG GGC TTG CTG GTG T |
| *FABP4* | ACG AGA GGA TGA TAA ACT GGT GG |
|  | GCG AAC TTC AGT CCA GGT CAA C |
| *MKI67* | GAA AGA GTG GCA ACC TGC CTT C |
|  | GCA CCA AGT TTT ACT ACA TCT GCC |
| *CD36* | CAG GTC AAC CTA TTG GTC AAG CC |
|  | GCC TTC TCA TCA CCA ATG GTC C |
| *GLUT4* | TGG GCT TCT TCA TCT TCA CC |
|  | GTG CTG GGT TTC ACC TCC T |
| *GCK* | CAT CTC CGA CTT CCT GGA CAA G |
|  | TGG TCC AGT TGA GAA GGA TGC C |
| *CPT1b* | TGT ATC GCC GTA AAC TGG ACC G |
|  | TGT CTG AGA GGT GCT GTA GCA C |
| *Gapdh* | TGC ACC ACC AAC TGC TTA G |
|  | GGA TGC AGG GAT GAT GTT C |
| *Ucp1* | GTG AAC CCG ACA ACT TCC GAA |
|  | TGA AAC TCC GGC TGA GAA GAT |
| *Ppargc1a* | ACA GCT TTC TGG GTG GAT TG |
|  | TGT CTC TGT GAG GAC CGC TA |
| *Prdm16* | TAT GGA GTG ACA TAG AGT GTG CT |
|  | CCA CTT CAA TCC ACC CAG AAA G |
| *Glut4* | GTA ACT TCA TTG TCG GCA TGG |
|  | AGC TGA GAT CTG GTC AAA CG |
| *Gck* | GCA TCT CTG ACT TCC TGG ACA AG |
|  | CTT GGT CCA GTT GAG CAG GAT G |
| *Cd36* | GGA CAT TGA GAT TCT TTT CCT CTG |
|  | GCA AAG GCA TTG GCT GGA AGA AC |
| *Cpt1b* | ATG TAT CGC CGC AAA CTG GAC C |
|  | CTC TGA GAG GTG CTG TAG CAA G |
| *Mki67* | GAG GAG AAA CGC CAA CCA AGA G |
|  | TTT GTC CTC GGT GGC GTT ATC C |
| *UCP1 gRNA-1* | CCCGAGGCACCGAGCGAGAAT |
| *UCP1 gRNA-2* | GCAGGGCTCCCGAGGCACCGA |
| *UCP1 gRNA-3* | AGGGCTCCCGAGGCACCGAGC |
| *UCP1 gRNA-4* | ACCGAGCGAGAATGGGAATGG |
| *UCP1 gRNA-5* | GCACCGAGCGAGAATGGGAAT |
| *PPARGC1a gRNA-1* | CACTGAAGCAGAGGGCTGCCT |
| *PPARGC1a gRNA-2* | TTAGAGCAGCAAGCTGCACAG |
| *PPARGC1a gRNA-3* | GTTAGAGCAGCAAGCTGCACA |
| *PPARGC1a gRNA-4* | GGGCTGCCTTTGAGTGACGTC |
| *PPARGC1a gRNA-5* | AGTTAGAGCAGCAAGCTGCAC |
| *PRDM16 gRNA-1* | CGGCGGCGGCGGCGCGACGAT |
| *PRDM16 gRNA-2* | GGCGGCGCGACGATGAGGATG |
| *PRDM16 gRNA-3* | GCCGCCGCCGCCGCCTCGGCG |
| *PRDM16 gRNA-4* | CGCCGCCGCCGCCGCCTCGGC |
| *PRDM16 gRNA-5* | GGCGGCGGCGGCGGCGCGACG |
| *Ucp1 gRNA-1* | GGGACTTGGCAGGGGCGTGCC |
| *Ucp1 gRNA-2* | CTGAGCCGGCCCAGGTCTCCA |
| *Ucp1 gRNA-3* | CAGGTCTCCAAAGAGCTGCTA |
| *Ucp1 gRNA-4* | CTTTGGGAGTGACGCGCGGCT |
| *Ucp1 gRNA-5* | GGCTTTGGGAGTGACGCGCGG |

**Table S3.** List of antibodies.

| Name | Vendor | Concentration | Catalog number |
| --- | --- | --- | --- |
| Ki67 (SolA15) | Fisher Scientific | 5ug/ml | 14-5698-82 |
| Carbonic Anhydrase | Fisher Scientific | 3ug/ml | AF2188 |
| CD31 | Fisher Scientific | 10ug/ml | BBA7 |
| Goat anti-rat, Alexa Flour 647 | Fisher Scientific | 1:1000 | A21247 |
| Goat anti-mouse, Alexa Fluor 594 | Life Technologies | 1:500 | A11032 |
| Donkey anti-goat, Alexa Fluor 594 | Fisher Scientific | 1:1000 | A11055 |
